# Coordination during group departures and group progressions in the tolerant multilevel society of wild Guinea baboons (*Papio papio*)

**DOI:** 10.1101/797761

**Authors:** Davide Montanari, Julien Hambuckers, Julia Fischer, Dietmar Zinner

## Abstract

**Objectives:** Most primate species live in groups, and temporal and spatial coordination of activities of individuals is essential for maintaining group cohesion, and there is still considerable debate to which degree social organization, the extent of despotism, and resource distribution shape group coordination processes. As different baboon species exhibit considerable variation in all of these factors, they constitute an excellent test case to resolve this debate.

**Materials and Methods:** We analyzed group departures and progressions of Guinea baboons, *Papio papio*, in the Niokolo Koba National Park in Senegal. Guinea baboons live in a multi-level society with strong male bonds and a lack of a clear dominance hierarchy between males.

**Results:** Two-thirds of departures were initiated by adult males, and one third by adult females. Although initiators were more likely to signal than followers, signaling did not affect the initiation success. During group progression, males that were not affiliated with females were predominantly found in the front, while affiliated males, females and young were observed more frequently closer to the center of the group, and no preferences for rear positions. Overall, affiliated subjects were more likely to depart and travel together.

**Discussion:** The group departures in Guinea baboons differed strikingly from the elaborate ‘negotiation’ behaviors among male hamadryas baboons, *Papio hamadryas*. We did not observe that specific individuals dominated the group coordination. Neither social organization, variation in despotism, nor resource distribution alone explain variation between species. Future studies should test whether specific combinations of factors promote the occurrence of negotiation processes.

**Research Highlights:**

- In wild Guinea baboons, both adult males and females initiated group departures
- Initiators signaled during departures, but this did not affect initiation success
- Solitary males were predominantly found at the front during group progression

## 1 INTRODUCTION

Taking advantage of the benefits of group living requires the temporal and spatial coordination of activities of individuals (Conradt & Roper, 2003; King & Cowlishaw, 2009; Krause & Ruxton, 2002). The coordination of individual movements in particular is essential for maintaining group cohesion (Couzin, Krause, Franks & Levin, 2005; Petit & Bon, 2010; Westley, Berdahl, Torney & Biro, 2018). These coordination processes differ between species and contexts. Many of the coordination patterns seen in swarms, flocks and certain social groups such as desert locusts, *Schistocerca gregaria* (Bazazi et al., 2008), European starlings, *Sturnus vulgaris* (Ballerini et al., 2008), three-spined sticklebacks, *Gasterosteus aculeatus* Jolles, Boogert, Sridhar, Couzin & Manica, 2017), but also some mammal species (e.g., herds of domestic sheep, *Ovis aries*, Toulet, Gautrais, Bon, & Peruani, 2015) can be explained by simple rules characterizing the attraction, alignment, repulsion and mimetism between neighboring individuals (Couzin & Krause, 2003; Deneubourg & Goss 1989; Sueur & Deneubourg, 2011). However, especially in socially complex societies, several other factors can modulate group coordination mechanisms and processes, such as individual traits (Couzin et al., 2011; del Mar Delgado et al., 2018), heterogeneous social relationships (Nagy, Ákos, Biro & Vicsek, 2010), or landscape features (Strandburg-Peshkin, Farine, Crofoot & Couzin, 2017).

To investigate how group coordination arises from individual decisions, most studies focused on the initiation of collective movements after the group had been stationary for a while (e.g., Kummer 1968a; Stueckle & Zinner, 2008). Key questions here are who attempts to initiate group movements and whether attempts are successful, i.e. whether other group members follow and in which order (e.g., Black, 1988; Lorenz, 1931; Stolba, 1979; Sueur & Petit, 2008a,b; Walker, King, Mcnutt & Jordan, 2017). In some species, the initiation of group movements is highly biased towards certain individuals, often dominant or old and experienced group members (e.g., mountain gorillas, *Gorilla beringei beringei*, Watts, 2000; bottlenose dolphins, *Tursiops* sp., Lusseau & Conradt, 2009). Such cases have been described as consistent leadership (Conradt & Roper, 2005; Pyritz, King, Sueur & Fichtel, 2011a; Strandburg-Peshkin, Papageorgiou, Crofoot & Farine, 2018). In other species, initiation attempts are distributed among many or all (often only adult) group members (e.g., meerkats, *Suricata suricata*, Bousquet, Sumpter & Manser, 2011; white-faced capuchins, *Cebus capucinus*, Leca, Gunst, Thierry & Petit, 2003). These cases have been characterized as distributed or variable leadership (Conradt & Roper, 2005; Pyritz et al., 2011a; Strandburg-Peshkin et al., 2018).

The propensity to initiate group movements can be affected by individual, social or environmental factors (Farine, Strandburg-Peshkin, Couzin, Berger-Wolf, & Crofoot, 2017). Initiators often belong to specific age and sex-classes. For instance, old female bonobos, *Pan paniscus* (Tokuyama & Furuichi, 2017) or adult female European bisons, *Bison bonasus* (Ramos, Manizan, Rodriguez, Kemp & Sueur, 2018) initiate the majority of movements. Individual physiological needs can also modulate the initiation process: lactating plain zebra females (*Equus burchellii*) initiate collective movements more frequently than non-lactating females (Fischhoff et al., 2007). Bolder individuals may initiate group movements more often than shy group members (e.g., domestic horses, *Equus ferus caballus*, Briard, Dorn, & Petit, 2015; red-fronted lemurs, *Eulemur rufifrons*, Sperber, 2018). In groups where power differentials play an important role, that is in more despotic societies, high ranking subjects are more likely to initiate group movement (e.g., despotic rhesus macaques, *Macaca mulatta*, than in more egalitarian Tonkean macaques, *Macaca tonkeana*, Sueur & Petit, 2008a). The social organization (uni-level vs. multi-level) is also expected to modulate group coordination processes (Fishhoff et al., 2007; Ozogány & Vicsek, 2015; Sueur et al., 2011). Finally, environmental heterogeneity could modulate the propensity to initiate a group movement as it modulates collective behavior in a variety of ways (e.g., Bonnell, Henzi & Barrett, 2019; King et al., 2018; Strandburg-Peshkin et al., 2017).

Baboons (genus *Papio*) are an intriguing model to study the impact of social factors on group coordination, as they exhibit considerable variation in mating system, social organization and social structure (Anandam et al., 2013; Fischer et al., 2017; Swedell, 2011). Olive (*P. anubis*), yellow (*P. cynocephalus*), chacma (*P. ursinus*) and Kinda baboons (*P. kindae*) usually live in a uni-level, multimale-multifemale group (Anandam et al., 2013). Their societies reveal a linear rank hierarchy, determined through agonistic interactions in males and inherited in females (Anandam et al., 2013; Barrett & Henzi, 2008; Swedell, 2011). Hamadryas (*P. hamadryas*) and Guinea baboons (*P. papio*), in contrast, live in multilevel societies based on monandric-polygynic reproductive units (one-male units or OMUs) at the base of the societies (Fischer et al., 2017; Goffe, Zinner & Fischer, 2016; Kummer, 1968a,b; Patzelt et al., 2014; Pines & Swedell, 2011; Schreier & Swedell, 2009). Thus, baboons provide a useful model to compare group coordination in uni-level and multi-level societies.

Studies of group coordination in uni-level baboon societies have shown heterogenous results. In some groups, dominant males predominantly initiated and directed troop movements (chacma baboons, Byrne, Whiten & Henzi 1990; Stoltz & Saayman, 1970; but see Buskirk, Buskirk & Hamilton, 1974). In a further study of chacma baboons, adult males were more likely to initiate group movements but the likelihood of being successful was similar for males and females (Stueckle & Zinner, 2008). However, when provided with incentives, the dominant male led groups to experimental food patches (King et al., 2008). In olive baboons at Gombe, the highest-ranking male was also more likely to determine the direction and timing of group movements than lower ranking subjects (Ransom, 1981), whereas in Queen Elizabeth National Park high ranking males often attempted to initiate a group movement, but they were only successful when old females followed him (Rowell, 1969). A similar impact of high-ranking females has been observed in yellow baboons in Mikumi National Park (Norton, 1986). A recent study in which olive baboons were tracked with a high-resolution global positioning system revealed a process of shared decision-making characterizing group movement. Rather than preferentially following dominant individuals, these baboons were more likely to follow when multiple initiators agreed (Strandburg-Peshkin, Farine, Couzin & Crofoot, 2015).

In hamadryas baboons, which live in a multi-level society (Grueter & Zinner, 2004; Kummer, 1968a), the reproductive males of the OMUs almost exclusively initiated group movements, while females had only a little impact on group coordination (Kummer, 1968a, 1995; Stolba, 1979). In subgroups of two OMUs, Kummer (1968a) described the decision making process as a “negotiation” among males with different roles, the initiator and the decider male (ID-system). Initiators moved away from the center of the band followed by their females. If another male (decider) from the band did not follow, the initiator moved back to the center. The ID-system was, however, not confirmed in a subsequent study on the same population, when larger social entities were taken into account (e.g. clans, bands; Stolba, 1979).

Guinea baboons live in a similar multi-level social organization as hamadryas baboons. If the social organization affects decision making, one could expect a similar strong impact of OMU males on the initiation of collective movements as in hamadryas baboons. However, Guinea baboon males are socially more tolerant than hamadryas baboon males and Guinea baboon females are not as strictly controlled by their males (Fischer et al., 2017; Kummer, 1968a), which might also affect the females’ role in initiation collective movements. Thus, if the socially more tolerant style modulates the decision-making process during group departures, one would expect that females take a share in the initiation of group movement.

In both types of baboon social organization, individuals appear to preferentially follow closely affiliated group members, irrespective of who initiates a group movement (olive baboons, Farine et al., 2016; chacma baboons, King et al., 2008, 2011). In hamadryas baboons the departure process relies on unit member cohesiveness (Kummer, 1968a, 1995). We therefore expected that the relationship strength affects who is likely to follow whom during group departures, with animals having stronger relationships being more likely to depart in close succession.

We additionally investigated the function of signals in group departures. Signals are conceived as indicators of specific behavioral dispositions (Fischer & Price, 2017). Thus, subjects who are motivated to initiate a group movement should express this motivation using signals (e.g., bonobos, *Pan paniscus*, Schamberg, Cheney & Seyfarth, 2017). We therefore predicted that subjects who initiated group departures were more likely to signal compared to individuals whom we classified as followers. We furthermore predicted that subjects who signaled may indicate a greater decisiveness to initiate group movement, and therefore might be more successful in recruiting followers.

In the second part of this study, we investigated progression order. We focused on situations when the baboons moved in more or less a single-file. The progression order has been regarded as an adaptation to predation risk (DeVore & Washburn, 1963; Rhine, 1975; Rhine, Forthman, Stillwell-Barne, Westlund & Westlund, 1981; Rhine, Bioland, & Lodwick, 1985). DeVore and Washburn (1963) reported a socio-spatial order in which the most vulnerable group members (adult females, juveniles and infants) took central positions close to the dominant adult males, whereas low-ranking adult males and older immature males occupied the more risk prone positions in the front and rear of the progression. However, this male-centered order was not observed in other baboon populations (Altmann, 1979; Rowell, 1969; Harding, 1977; Rhine, 1975; Rhine & Westlund, 1981; Rhine et al., 1985; Rhine & Tilson, 1987). For multi-level hamadryas baboons, Kummer (1968a) reported that the frequency with which adult and subadult males appeared at the front was twice that which would be expected by chance, whereas males were found at the rear with a frequency equal to chance.

Regarding group progressions, we therefore contrasted two possible scenarios: if Guinea baboons conform to other baboon species, adult males should be found more frequently in front and rear positions, while adult females and youngsters should mainly travel in the middle of the progression. Alternatively, units may retain their cohesiveness during group movement. In this case, the progression would resemble the male centered pattern with primary unit males moving with their females and offspring in the center of a progression.

## 2 MATERIALS AND METHODS

### 2.1 Field site and study subjects

The fieldwork was conducted in the surroundings of the field station “Centre de Recherche de Primatologie (CRP) Simenti” (13°01’34” N, 13°17’41” W), in the Niokolo-Koba National Park, south-eastern Senegal. The multi-level system of Guinea baboons consists of “units” (usually one adult male and one to several females with their young), units are nested within “parties” and parties are nested within “gangs” (Fischer et al., 2017). The study subjects were fully habituated baboons belonging to five parties, that formed two gangs (Table 1). Subjects were individually identified, although the identification of juveniles was not always possible. The home ranges of the parties covered on average 30.3 km^2^ of largely overlapping territories (Kernel density estimations 95%, unpubl. data, M. Klapproth).

**Table 1.** Average composition of study groups. Party sizes (i.e. total number of party members) varied due to births, deaths, disappearances, between-parties transfers of individuals and difficulties in recognizing young weaned individuals.

| Gang | Party | Number of units | Number of adults |  | Size |
| --- | --- | --- | --- | --- | --- |
| "Mare" | "4" | 2-3 | 5 ♂ | 3 ♀ | 15 |
|  | "9" | 5-6 | 12 ♂ | 17 ♀ | 45 |
|  | "10" | 1-2 | 2 ♂ | 2 ♀ | 8 |
| "Simenti" | "5" | 3-4 | 10 ♂ | 9 ♀ | 25 |
|  | "6" | 4-5 | 12 ♂ | 11 ♀ | 38 |

### 2.2 Data collection

Data collection was conducted from January to August, in 2016 and 2017, for a total of 16 months, 6 days per week. Observation days started before sunrise (at 6:00 or 6:30) to locate the baboons at the sleeping site. Data were recorded on Samsung Note 3 handhelds using forms created with Pendragon 7.2 (Pendragon Software Corporation, USA). Every day, all researchers working at CRP collect census, ad libitum, scan, and focal data of the baboons to investigate the demography, reproductive success, association data, and behavioral patterns (Altmann, 1974). These data were used to determine female-male associations. Data on group movement were collected with the all-occurrence sampling method (Altmann, 1974). Two types of events were distinguished during the group movement process: group departures and group progressions (see below). We classified individuals according to age (Category “young” including infants, yearlings, and juveniles; Category “adult” including subadults and adults) and sex. We further noted the unit identity for primary males and the associated members of the unit. Non-primary adult males (i.e. secondary and unaffiliated ones) and young individuals which could not be unambiguously identified as members of one unit were labelled by their own IDs. In addition we considered the unit size (number of adult subjects). Non-associated animals had a unit size of 1, units comprised of an adult male and one female had a size of 2, and so on. The largest unit size was 7.

#### 2.2.1 Operational definition of group departures

A group departure occurred when a group of baboons was collectively leaving a confined area where they had been stationary for a set time. We collected data on events of group departures throughout the day, whenever visibility allowed it and certain conditions were met. Specifically, the group had to consist of one or more complete units or a complete party. The confined area where the individuals stayed stationary before a group departure was named the pre-departure area. The size of the pre-departure area was 20 m in diameter at maximum. The individuals had to be isolated from conspecifics outside the area for at least 20 m. The individuals had to stay stationary, either feeding, resting, or socializing in the pre-departure area for at least 15 minutes, to ensure a certain degree of independence in timing and direction from previous movements (comparably to e.g., Leca et al., 2003; Pyritz, Kappeler & Fichtel 2011b; Seltmann, Majolo, Schülke & Ostner, 2013; Sueur & Petit, 2008a,b). We excluded movements prompted by predation risks, alarm calls or social interactions such as threats or chases. When these conditions were met, the identity of all individuals moving away from the pre-departure area and the starting time and the direction of their movements were voice recorded.

The first individual leaving the area was defined as attempting an initiation of group departure. The individuals moving away from the pre-departure area in the same direction as another one before, within a 5-minute interval time, were considered followers. When an individual was heading more than 45° to the left or right from the direction chosen by the previous individual, and/or was starting to move away more than 5 minutes after the previous individuals, it was coded as attempting another initiation of group departure. Therefore, an initiation attempt was coded as successful when some or all individuals in the pre-departure area followed. All individuals of the subject group were hence classified as successful initiators, unsuccessful initiators, or followers. Unsuccessful initiators were subsequently coded either as followers, successful initiators or again as unsuccessful initiators on the following initiation attempt. When two successful initiations were coded in one event, this implied group fission.

We furthermore recorded whether any one of the following signals occurred, to test whether they signaled the readiness to initiate a group departure or affected the likelihood to succeed in initiation:

- Back glance: once the individual has started to move away from the pre-departure area and it looks back in the direction of other group members. Empirically defined as the turn of the head of more than 90° towards the direction of the pre-departure area.
- Branch shaking display: rapid repeated bouncing in place while the individual stands quadrupedal grasping a flexible branch, shaking it (Mehlman, 1996).
- Pause: once the individual has started to move away from the pre-departure area and it stops moving for more than 2 seconds within the first 20 m of movement.
- Vocalizations: individual call, classified per type: keck, grunt, roar grunt, scream, bark, wahoo (Fischer et al., 2017; Maciej, 2013).
- Greeting: “exchange of non-aggressive signals that consist of species-specific behavioral patterns, […] ranging from touches and embraces to genital manipulation and same-sex mounts” (Dal Pesco & Fischer, 2018, p. 88).

#### 2.2.2 Operational definition of group progressions

A group progression was defined as the instance when a group of travelling baboons was positioned in an approximate single-file and jointly moved in (largely) the same direction. Single-file travel progressions typically occur along delineated pathways such as roads and on open areas. We collected data on events of group progressions throughout the day, whenever visibility allowed it and the following conditions were met: the progressing group had to consist of one or more complete parties and the first data regarding a group progression event had been collected at least 30 minutes after the end of a previous event. When these conditions were met, D.M. advanced a few meters in front of the moving group, stopped and set a virtual reference line on the ground in front of the arriving group. Whenever a baboon crossed this reference line, its identity and time of crossing (to the nearest second) were voice recorded.

### 2.3 Data analyses

All models and plots were fitted in R (version 3.5.0; R Core Team, 2018), using RStudio interface (version 1.1.383; RStudio Team, 2016). The only exception concerns the representation of posterior probability distributions of the order of group progression. These plots were created with MATLAB (version 9.4; The MathWorks, Inc., 2018).

#### 2.3.1 Group departures

We first tested whether the likelihood of attempting an initiation of group departure was influenced by sex, age and/or unit size. To this end, we ran a Generalized Linear Mixed Model (GLMM; Baayen, 2008) with a binomial response variable and logit link function. Sex, age and unit size were included as fixed effects, individual identity and event as random effects (both random intercept components) and time of the day as a polynomial predictor variable. To prevent any scaling issue, we applied a z-transformation of the time of the day. We used the function glmer provided by the R package lme4 (version 1.1-17; Bates, Mächler, Bolker & Walker, 2015), setting the optimizer to ‘bobyqa’ to prevent convergence issues. To test if the full model fits better than a simpler alternative with a likelihood ratio test (Dobson, 2002), we compared the full model to the null model containing only the random effects and time. The p-values for the distinct effects were derived comparing the full model with the model reduced of the predictor of interest, using the function drop1, argument ‘test’ set to ‘Chisq’. To obtain the confidence intervals for the different regression coefficients, we used a bootstrap procedure using the function bootMer provided by lme4 (nboots = 1000). In a second step, with the same procedure, we tested whether the same set of independent variables was affecting the success of the initiation attempts.

In order to approximate distances between individuals and to investigate the individual spatial association within the party, we calculated interval times (to the nearest second) between dyads of individuals succeeding each other. We restricted the analysis to those 40 events where at least one complete party was present, and calculated interval times only for individually identified subjects (omitting most of the juveniles).

To test whether interval times were influenced by unit identity, we used a linear mixed model (LMM; Baayen, 2008) into which we included unit membership, that is, whether individuals belonged to the same unit as fixed effect, and the identity of the individual following, i.e. for which we calculated the interval time, as well as the event as random effects. The model was fitted using the function lmer of the R package lme4 (version 1.1-17; Bates et al., 2015). Because the interval times were highly skewed, they were log-transformed. We verified that the assumptions of normally distributed and homogeneous residuals were met by visually inspecting a qqplot and a plot of the residuals against the fitted values. Both plots indicated that the assumptions were met. We tested model stability by excluding subjects one by one from the dataset and comparing the model estimate outcomes of these subsets with those outcomes of the full dataset. This revealed no influential subjects. We tested whether the full model was significantly better compared to the null model, in which the fixed effect was omitted, with the R function anova (argument test ‘Chisq’; Dobson, 2002; Forstmeier & Schielzeth, 2011). The models were fitted using Maximum Likelihood, rather than Restricted Maximum Likelihood, to allow for a likelihood ratio test (Bolker et al., 2009). The p-value for the fixed effect was based on a likelihood ratio test comparing the full with the reduced model, with the function drop1, argument ‘test’ set to ‘Chisq’ (Barr, Levy, Scheepers & Tily, 2013).

#### 2.3.2 Group progressions

To test whether specific individuals would be preferentially found in specific parts of the group, we divided the sequence of individuals into equal thirds. We used a multinomial logit regression model with random intercepts (Fahrmeir, Kneib, Lang & Marx, 2013). Progression-location was coded into three categories (front, middle and rear), with the probability of belonging to the category conditioned on age (adult vs young) and on one variable with three terms: female, primary male, non-primary male (“f_pm_npm”). The model was estimated by means of Bayesian methods. Posterior densities of the regression coefficients were obtained from Markov-chain Monte Carlo (MCMC) procedures, using the R package MCMCglmm (Hadfield, 2010). From the resulting posterior samples of progression-location regression coefficients, we calculated the distribution of the relative frequency (i.e. the probability p) to observe a progression-location k = 1, 2, 3, conditional on age = adult (ESM formula set 1), as well as the distribution of the relative frequency to observe progression-location k = 1, 2, 3, conditional on f_pm_npm = female (ESM formula set 2).

In addition, we ran a post-hoc test to investigate whether non-primary males were occupying edge positions during group progressions compared to primary males. To do this, we divided the sequence of individuals of the front third and the one of the rear third in two equal parts. We ran a GLMM with a binomial response variable and logit link function. We used the function glmer provided by the R package lme4 (version 1.1-17; Bates et al., 2015). f_pm_npm was introduced as one fixed effect with three levels: female, primary male, non-primary male. Individual identity was included as a random effect. Model diagnostics were performed by creating scaled residuals through simulations from the fitted model with the function simulateResiduals (number of simulations: 1000), provided by the R package DHARMa (version 0.2.0; Hartig, 2017). We also plotted the residuals against the predicted response from the model, using the function plotSimulatedResiduals, provided by the R package DHARMa. The plot permits to detect deviations from uniformity in y-axis direction and performs a quantile regression, which provides 0.25, 0.50 and 0.75 quantile lines across the plots. Reported p-values for the individual effects were obtained from likelihood ratio tests comparing the full with the respective reduced models (R function drop1, Barr et al., 2013).

Finally, we investigated the spatial association within the progressing party to test whether interval times were influenced by unit membership, as for group departures. We measured the time differences between individuals to the nearest second and used the same procedure applied to the dataset of group departures. In brief, we used a linear mixed model (LMM; Baayen, 2008) into which we included unit membership, that is, whether individuals belonged to the same unit as a fixed effect, and the identity of the individual following, i.e. for which we calculated the interval time, as well as the event as random effects.

## 3 RESULTS

### 3.1 Group departures

We collected data during 121 group departure events. Thirty-three events involved only one complete unit, 48 events involved more than one complete unit, and 40 events involved a complete party. In total, we sampled 146 attempts of group departure: 52 (35.6%) conducted by adult females, 91 (62.3%) by adult males and 3 (2.1%) by juveniles. Twenty-three attempts of initiation were not successful (15.8%) (Table 2). In two events, the individuals in the departure area split during group departure, after two successful initiation attempts within the same event. Fifty-eight different individuals attempted to initiate a group departure: 28 different adult males, 27 different adult females and three different juveniles.

**Table 2.** Number of initiation attempts by adult females, adult males, and young in relation to the level of social organization and initiation success.

| Level of social organization | initiation | adult female | adult male | young |
| --- | --- | --- | --- | --- |
| one unit | successful | 15 | 16 | 2 |
|  | unsuccessful | 4 | 1 | 0 |
| more units | successful | 18 | 32 | 0 |
|  | unsuccessful | 2 | 6 | 1 |
| party | successful | 9 | 31 | 0 |
|  | unsuccessful | 4 | 5 | 0 |

Overall, the predictors age and sex had a clear impact on the probability of attempting an initiation of group departure (likelihood ratio test comparing full and null model: χ^2^ = 71.882, df = 6, P < 0.001). Being male and of adult age strongly increased the likelihood of attempting an initiation. Within the different adult age categories, there was no difference in the likelihood to initiate a group departure (Table 3, Figure 1).

**Table 3.** Effects of age and sex category, as well as unit size, and time of day on the likelihood of attempting to initiate a group departure. Estimated coefficients, standard errors, confidence intervals, and test statistics.

| | Estimate | Std. Error | CI<sub>lower</sub> | CI<sub>upper</sub> | $\chi^2$ | Df | P |
| --- | --- | --- | --- | --- | --- | --- | --- |
| Intercept | -6.113 | 0.793 | -7.385 | -4.898 | (1) | (1) | (1) |
| sex Male | 1.047 | 0.231 | 0.625 | 1.533 | 14.865 | 1 | <0.001 |
| age Mature adult | 3.643 | 0.726 | 2.551 | 4.545 | 66.680<sup>(2)</sup> | 4 | <0.001<sup>(2)</sup> |
| age Old adult | 3.878 | 0.766 | 2.619 | 5.022 | (2) | (2) | (2) |
| age Subadult | 3.591 | 0.764 | 2.387 | 4.734 | (2) | (2) | (2) |
| age Young adult | 3.399 | 0.745 | 2.249 | 4.389 | (2) | (2) | (2) |
| unit size | 0.028 | 0.078 | -0.126 | 0.183 | 0.122 | 1 | 0.727 |
| z.time | 0.104 | 0.101 | -0.108 | 0.317 | (1) | (1) | (1) |
| l(z.time^2) | -0.057 | 0.063 | -0.237 | 0.052 | 0.877 | 1 | 0.349 |

<sup>(1)</sup> not meaningful in this context; <sup>(2)</sup> equal values because they refer to different terms of the same variable

**Figure 1.**
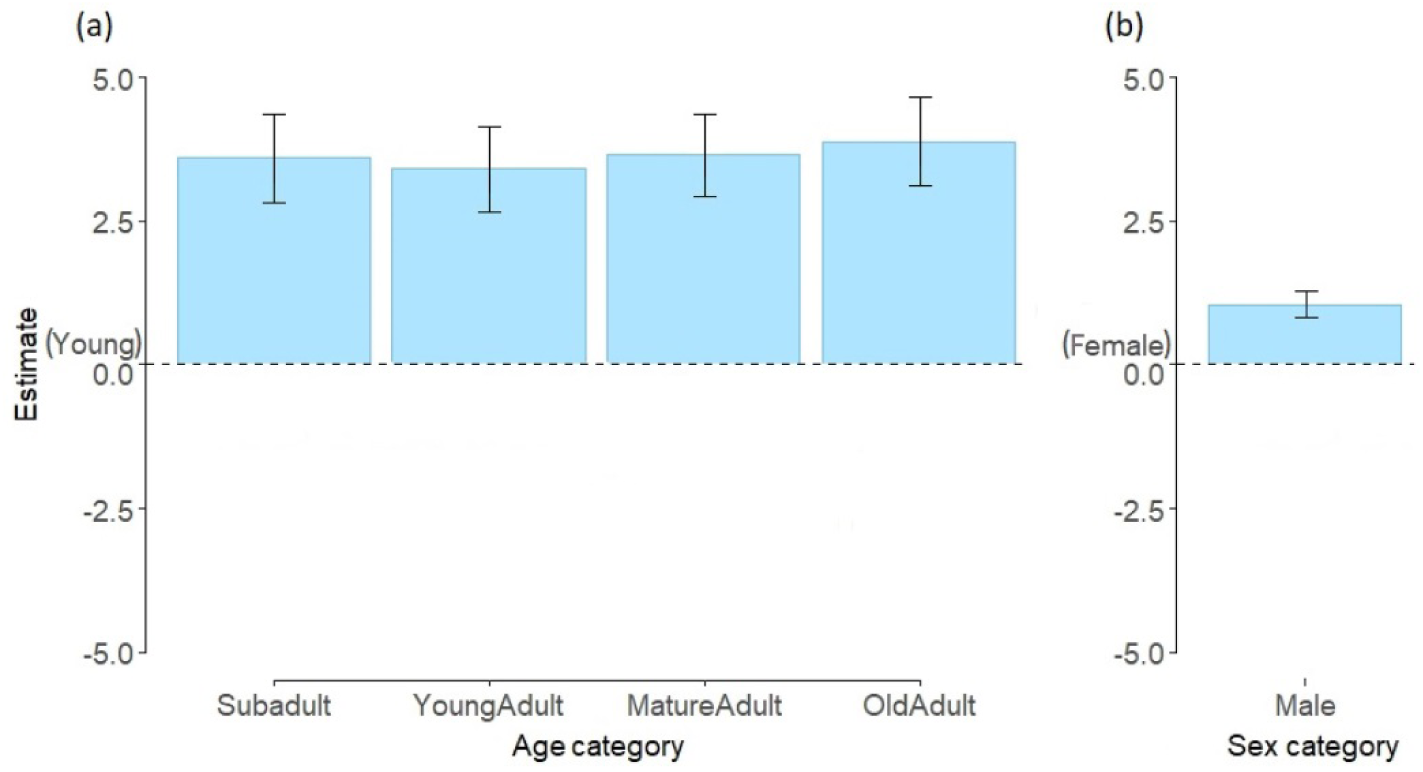
Estimates of the predictors age and sex on the likelihood of attempting an initiation of group departure, from GLMM (reference category being “young” and “female”). (a) All adult categories are significantly more likely to attempt than young individuals. (b) Males are significantly more likely to attempt than females.

Because only three group departures were initiated by young subjects, we excluded these from further analyses to avoid convergence issues. Out of the 52 initiation attempts by adult females, 42 (80.8%) were successful, while out of the 91 attempts by adult males, 79 (86.8%) were successful. Once failed, an individual that attempted to initiate tried again only twice in 23 occurrences of unsuccessful attempts. Adult age category, sex, and/or individual association did not explain the variation in success of initiation (likelihood ratio test: χ^2^ = 3.309, df = 5, P = 0.653).

We next tested whether initiators and followers differed in signal usage during group departures. Initiators signaled in 57.7% of observations, while followers used signals only in 19.5 % of observations (Figure 2a; mean signaling rates across N = 86 individuals; N = 1102 events; P < 0.001; Table S1). Whether or not initiators used signals had no effect on their success rates. When a signal was used, the success rate was 83.6%; when no signal was used, it was 86.4 % (Figure 2b; N = 142 events; P = 0.947, see ESM for details Table S2). Note that signaling rates were first averaged for each individual and then across all individuals.

**Figure 2.**
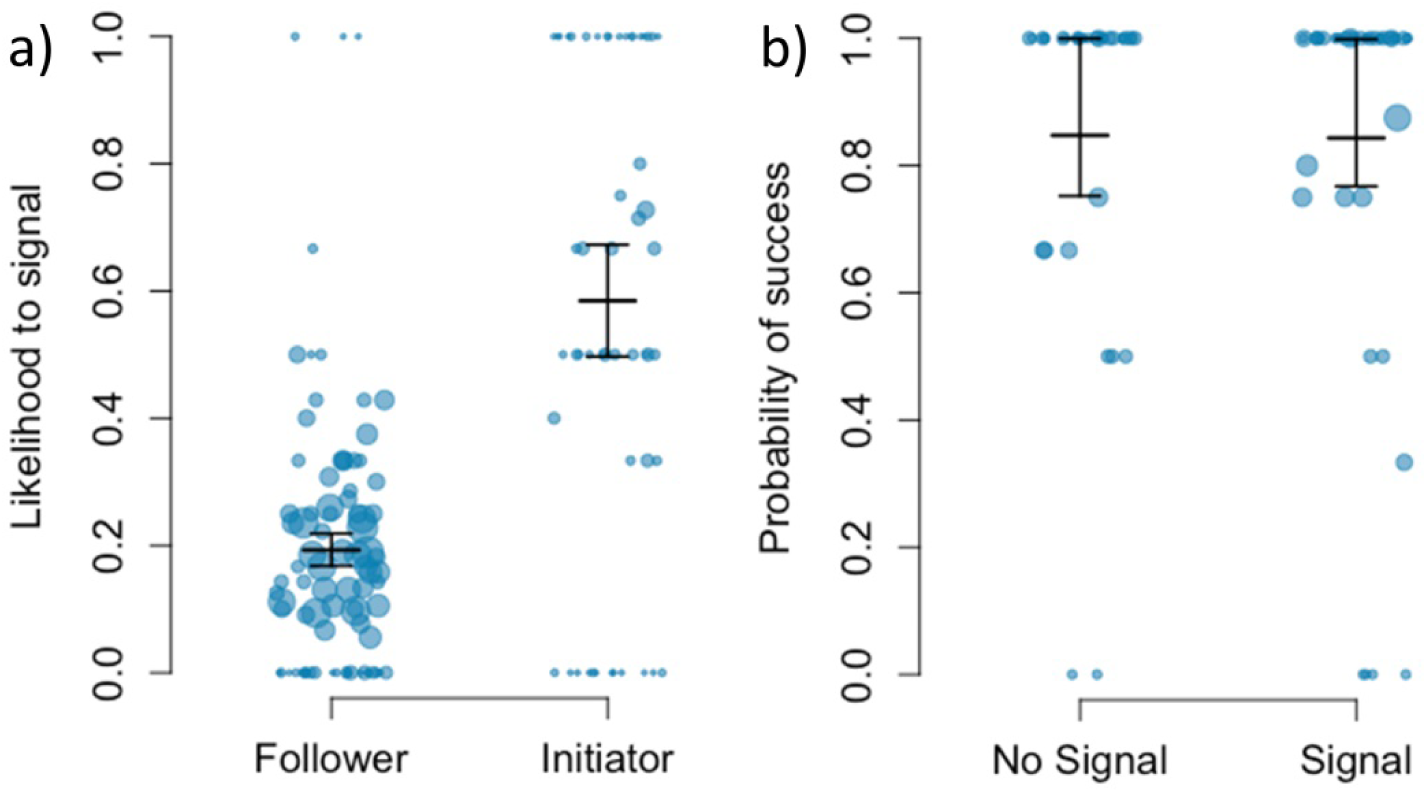
(a) Likelihood of followers and initiators to signal during group departures. (b) Probability of success in relation to signaling during initiation.

When leaving the pre-departure area, the time intervals between two individuals that belonged to the same unit was significantly shorter (mean = 13.7 s; range: 0-260 s) than the interval time between two individuals who did not belong to the same unit (mean = 25.6 s; range: 0-910 s); Table S3; likelihood ratio test: χ^2^ = 23.9, df = 1, P < 0.001, N = 813 intervals in 40 events).

### 3.2 Group progressions

We collected data on 100 events of group progression. Seventeen events involved more than one party. During the collected events, members of party 4 and 10 were always travelling with at least one of the other three parties. The number of events in which parties 4 and 10 were involved was very low (≤7 per party) compared to those in which party 5, 6 and 9 were involved (≥27 per party). Therefore, we excluded the individuals belonging to party 4 and 10 from the analyses, to achieve comparable numbers of events per party. Eleven events involved portions of a party because the party split for some hours or the whole day. In 6 of these events, the progressing group consisted of only 2 units.

Overall, the model outcomes revealed that age explained parts of the positioning of individuals during group progressions (i.e. 95% posterior density intervals do not include 0; Table 4). Adults were located more in front positions than middle or rear. It was also more likely to find adults in rear positions than in the middle of the group. Young individuals were somewhat less likely to take front positions compared to the other two categories (Figure 3a; the distribution of relative frequencies in Table S4).

**Table 4.** Effect of age (adult; young) on the likelihood for an individual to take a front, middle or rear position during a group progression. Reference category front and adult. Posterior means, confidence intervals, sample size and P-values derived from MCMC procedure.

|  | Posterior mean | CI<sub>lower</sub> | CI<sub>upper</sub> | effective sample size | P MCMC |
| --- | --- | --- | --- | --- | --- |
| middle and adult | -0.338 | -0.526 | -0.141 | 538.0 | <0.001 |
| rear and adult | -0.247 | -0.436 | -0.054 | 574.7 | 0.001 |
| middle and young | 0.542 | 0.177 | 0.886 | 648.9 | 0.004 |
| rear and young | 0.430 | 0.085 | 0.759 | 801.0 | 0.016 |

**Figure 3.**
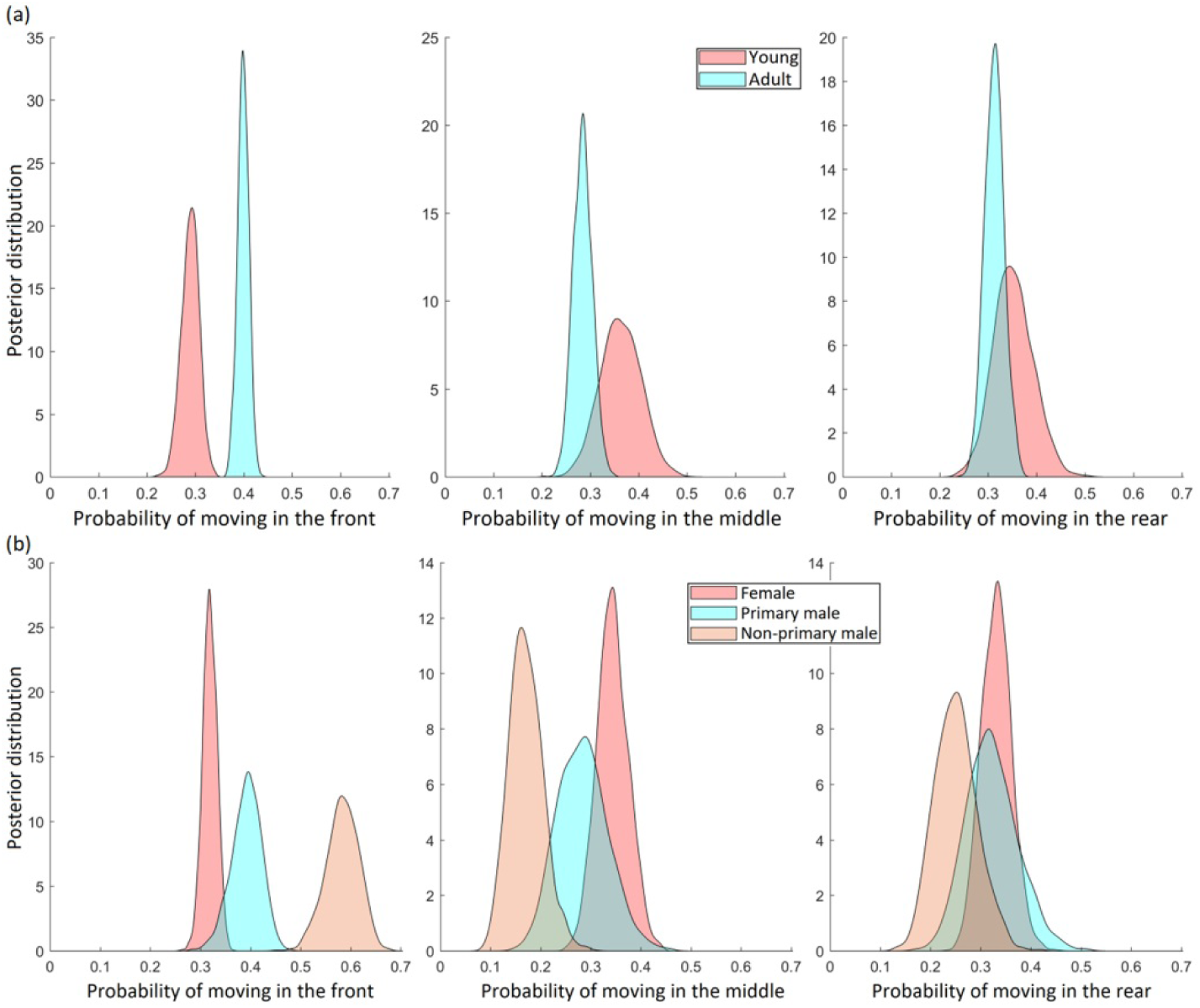
Posterior probability distributions to progress in front, middle or rear positions according to (a) age and (b) sex (adult subjects only). The distribution of relative frequency per category per third, i.e. estimated probabilities, in ESM.

We then considered only adult individuals for testing the effect of being a female, a primary male or a non-primary male on the position during group progressions. Sex and the distinction between primary and non-primary males explained variability in the order of group progression (Table 5). Adult females were found in all thirds with similar likelihood. Primary males mainly took front positions during group progressions, and were least frequently observed in middle positions. The strongest effects were observed for non-primary males, who were more likely to move in the front third than in the middle or rear third; their pattern differed significantly from that of females (distributions did not overlap; Figure 3b; the distribution of relative frequencies in Table S5)

**Table 5.**
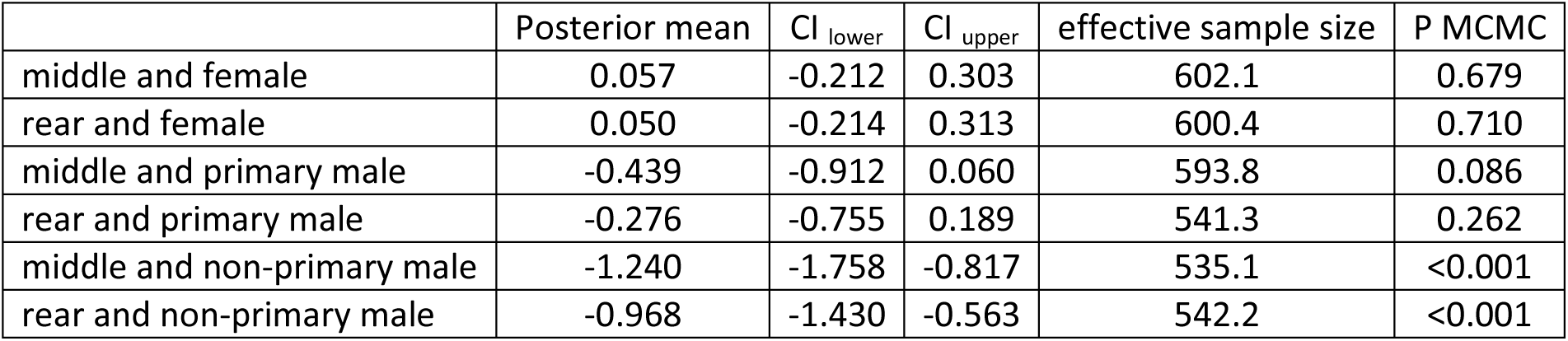
Effect of being a female, a primary male or a non-primary male on the likelihood for an individual to take front, middle or rear positions during a group progression. Reference category front third and female. Posterior means, confidence intervals, sample size and P-values derived from MCMC procedure.

Furthermore, non-primary males were observed significantly more often in the front half of the first third, as compared to females and primary males, which tended to progress in the half closer to the middle of the group (P < 0.001, Table S6). Non-primary males were also observed significantly more often in the back half of the rear third, as compared to females and primary males, which again progressed in the half closer to the middle of the group (P < 0.001, Table S7).

During group progressions subjects who belonged to the same unit were more likely to travel together, as evidenced by the interval time between two individuals belonging to the same unit (mean = 4.2 s; range: 1-70 s), which was significantly shorter than the interval time between two individuals that did not belong to the same unit (mean = 8.9 s; range: 1-293 s; likelihood ratio test: χ^2^= 201.5, df = 1, P < 0.001, N = 2226 intervals involving N = 120 individuals following in 100 events, Table S8).

## 4 Discussion

In our study population of Guinea baboons, collective movements were predominantly initiated by adult individuals. Adult males attempted initiations more often (62% of events) than adult females (36%, juveniles 2%). The vast majority of initiation attempts were successful (males 87%; females 80%). In other baboon species (olive, yellow and chacma), adult males were also reported as the major, but not exclusive, actors during group departures (King et al., 2011; Norton, 1986; Ransom, 1981; Stueckle & Zinner, 2008). The patterns we observed in Guinea baboon group departures and progressions were overall more similar to the patterns observed in uni-level species, such as chacma and olive baboons (Strandburg-Peshkin et al., 2015; Stueckle & Zinner, 2008), than to the patterns observed for hamadryas baboons.

In hamadryas baboons, only adult males were observed to take part in the negotiation and decision making on the direction and timing of coordinated departures of several OMUs (clans) from the sleeping sites (Kummer, 1968a, 1995; Stolba, 1979). In Guinea baboons, in contrast, adult females initiated group departures in about a third of the cases. Their greater share in initiating departures compared to other hamadryas baboons may be a result of the higher degree of ‘female freedom’. More specifically, female Guinea baboons are not coerced to maintain constant close proximity to their males and they have greater leverage in association patterns (Goffe et al., 2016). Also, the complex “negotiating” behaviors described for hamadryas baboons were observed extremely rarely. Instead, any adult Guinea baboon who moved off could trigger a group departure.

The observed differences between Guinea and hamadryas baboons likely reflect true species differences, but they may also be due to differences in data collection procedures. Descriptions of the hamadryas group departures by Kummer (1968a) and Stolba (1979) encompassed only departures from the sleeping site in a relatively open landscape, whereas our observations encompassed a mixture of observations in the early morning hours up to midday. We did not find any differences in departure processes among early morning departures and departures later during the day.

Although Byrne (1981) had observed negotiation processes similar to those described for hamadryas baboons during morning departures of Guinea baboons, we recorded such behaviors only in two cases. Males of two OMUs showed greeting interactions (Dal Pesco & Fischer 2018) before both left the sleeping site in the same direction with their party members. We are therefore rather confident that elaborate negotiation processes do not play a major role in group coordination in this species.

Another reason for the differences between Guinea and hamadryas in pre-departure coordination processes may be different ecological conditions of the two species (e.g. Chala, Roos, Svenning & Zinner, 2019). Kummer (1968a) and Stolba (1979) speculated that the elaborate coordination process of hamadryas baboons is an adaptation to their arid environment. To exploit food resources hamadryas bands often need to fission. Bands may break up into clans and even single OMUs during foraging, but have to fuse again at scarce water sources or sleeping sites. Since habitats of Guinea baboons in most parts of their distribution range are more productive than the average hamadryas baboon habitat, i.e. higher densities of food and water resources, an elaborate decision process on the direction of the daily travel direction might not be necessary.

Although signalers were more likely to use signals during departures, which could be taken as an expression of their intention to move (or perhaps their intention to initiate a group movement; Fischer & Zinner, 2011), this had no significant effect on their success in initiating group movement. However, the power to detect an effect of signaling was low, as initiators were generally highly successful in initiating group movement. It might also be the case that initiators who signaled were indeed more highly motivated than those who did not signal, while followers were not affected by the initiator’s expression of motivation (Fischer & Price 2017).

The spatial positioning of progressing baboons has been primarily seen as an adaptation to terrestrial lifestyle with its respective predation pressure (DeVore & Washburn, 1963). Progressions of olive baboons were led by low-ranking adult males and older immature males. The most dominant adult males, females with infants, and the youngest juveniles were in the center of the troop. The rear portion of the troop was a mirror image of the front, with low-ranking adult males and older immature males (DeVore & Washburn, 1963). In other populations of olive, yellow and chacma baboons, however, adult males predominantly occupied front positions, while young individuals mainly occupied central positions and adult females were equally spread from the front to the rear (Harding 1977; Rhine et al., 1985; Rhine & Tilson, 1987). The progression of Guinea baboons resembled the pattern described by DeVore & Washburn (1963), with non-primary adult males at the front and primary males in more central positions. Adult females, however, occupied front, center or rear position with similar probabilities, similar to what Rhine (1975) and Rhine & Tilson 1987) reported from yellow and chacma baboons. Positions at the rear of the group were equally taken by individuals of all age/sex classes. In summary, no clear pattern emerged for the different baboon species. The analysis of the interval times indicated that individuals belonging to the same units, i.e. individuals with closer social bonds, were more likely to depart and travel in close proximity, corroborating previous findings in other baboon species (Bonnell, Clarke, Henzi & Barrett, 2017; Farine et al., 2017; King et al., 2008, 2011; Kummer 1968a).

A comparison of the available data for the different species suggests that neither social organization nor ecological conditions fully account for differences in group coordination processes. With regard to the social organization, we found substantial differences between hamadryas and Guinea baboons; thus life in a multi-level society does not necessarily give rise to elaborate negotiation processes. The alternative idea that the harsh semi-desert conditions promotes negotiation behaviors and accounts for the observed variation neither seems to be true, as chacma baboons living in the Namib desert do not conform to the hamadryas pattern either (King et al., 2008, 2011). A possible explanation may be that it takes both factors together: a multi-level society with rather shallow rank hierarchies between males, and a resource distribution promoting fission-fusion dynamics. One way to test this conjecture would be to observe Guinea baboons living in harsh environments, such as the Sahara desert in Mauritania. Such observations are presently beyond our means, but could provide the answer to the question which combination of drivers accounts for the regulation of group coordination processes in baboons.

## Supporting information

ESM

## ACKNOWLEDGMENTS

We would like to thanks the Direction des Parcs Nationaux (DNP) and the Ministère de l’Environnement et de la Protection de la Nature (MEPN) du Sénégal for approval to conduct this study in the Parc National du Niokolo Koba (PNNK). The support of Col. Mallé Guye is particularly appreciated. We are grateful to the CPR team: Sonia Domínguez Alba, Muriel Drouglazet, Lauriane Faraut, Josefine Kalbitz, Harry Siviter, Franziska Wegdell for their help with the data collection and Roger Mundry for helpful discussion and statistical advice. This research was funded by the Deutsche Forschungsgemeinschaft (DFG, German Research Foundation) – Project number 254142454/GRK 2070, as part of the Research Training Group “Understanding Social Relationships”. JH was funded by the DFG RTG 1644 “Scaling problems in Statistics”. Support by the Leibniz ScienceCampus “Primate Cognition” is gratefully acknowledged.

## DATA AVAILABILITY STATEMENT

Data and code are available upon request.

