## Supplementary material for "Coordination during group departures and group progressions in the tolerant multilevel society of wild Guinea baboons (*Papio papio*)": ESM

**ELECTRONIC SUPPLEMENTARY MATERIAL**

### Supplementary Methods

**Formula set 1**: Distribution of the relative frequency (i.e. a probability denoted with p) to observe a progression-location k = 1, 2, 3, conditional on age = adult

$$\mathrm{P}_s(\text{progression} - \text{location} = k|\text{age} = adult) = \frac{\exp(\hat{\beta}_{adult,k})}{\sum_{j=1}^{3}\exp(\hat{\beta}_{adult,j})},$$

where adult is the reference category of the binary age factor, and as:

$$\mathrm{P}_s(\text{progression} - \text{location} = k|\text{age} = young) = \frac{\exp(\hat{\beta}_{adult,k} + \hat{\beta}_{young,k})}{\sum_{j=1}^{3}\exp(\hat{\beta}_{adult,j} + \hat{\beta}_{young,j})},$$

for age = young, where coefficient $\beta_{young,k}$ is the coefficient for changing the age factor from adult to young.

**Formula set 2**: Distribution of relative frequency to observe progression-location k = 1, 2, 3, conditional on f_pm_npm = female

$$\mathrm{P}_s(\text{progression} - \text{location} = k|\text{f\_pm\_npm} = female) = \frac{\exp(\hat{\beta}_{female,k})}{\sum_{j=1}^{3}\exp(\hat{\beta}_{female,j})},$$

where female is the reference category of the f_pm_npm factor, and as:

$$\mathrm{P}_s(\text{progression} - \text{location} = k|\text{f\_pm\_npm} = primary\ male) = \frac{\exp(\hat{\beta}_{female,k} + \hat{\beta}_{primary\ male,k})}{\sum_{j=1}^{3}\exp(\hat{\beta}_{female,j} + \hat{\beta}_{primary\ male,j})},$$

for f_pm_npm = primary male, where coefficient $\beta_{primary\ male,k}$ is the coefficient for changing the f_pm_npm factor from female to primary male, and as:

$$\mathrm{P}_s(\text{progression} - \text{location} = k|\text{f\_pm\_npm} = nonprimary\ male) = \frac{\exp(\hat{\beta}_{female,k} + \hat{\beta}_{nonprimary\ male,k})}{\sum_{j=1}^{3}\exp(\hat{\beta}_{female,j} + \hat{\beta}_{nonprimary\ male,j})},$$

for f_pm_npm = non-primary male, where coefficient $\beta_{nonprimary\ male,k}$ is the coefficient for changing the f_pm_npm factor from female to non-primary male. Index s = 1, …, S denotes the sample from the posterior density. To ensure the identification of the model, one progression-location category k ∈ {1, 2, 3} has to be selected as a reference category (Fahrmeir, Kneib, Lang, & Marx, 2013), for which we selected the front. For the Bayesian model fitting algorithm, all prior distribution assumptions were kept unchanged with respect to default options (namely Gaussian and inverse Gamma priors). MCMC sampling from the posterior distribution was performed for 40,000 iterations, with 15,000 burn-in iterations and a thinning by each 10$^{th}$ iteration. Therefore, the final posterior is based on 2,500 draws. We extracted posterior densities for the probability to belong to each third, for a given age and/or f_pm_npm class, and high posterior density intervals to further describe the effect of age and f_pm_npm on the probability to belong to a given progression-location category.

From the resulting posterior samples of progression-location regression coefficients (column "Posterior mean"), the distribution of relative frequency (i.e. estimated probabilities, Table S4), was calculated following the formulas explained. For example:

Probability for an adult to progress in the front

$$\frac{\exp\left(\hat{\beta}_{adult,k}\right)}{\sum_{j=1}^{3}\exp\left(\hat{\beta}_{adult,j}\right)} = \frac{1}{(1+\exp(-0.338)+\exp(-0.247))} = 0.401,$$

or, probability for a young individual to progress in the back

$$\frac{\exp\left(\hat{\beta}_{adult,k}+\hat{\beta}_{young,k}\right)}{\sum_{j=1}^{3}\exp\left(\hat{\beta}_{adult,j}+\hat{\beta}_{young,j}\right)} = \frac{\exp(-0.247)+\exp(0.430))}{(1+\exp(-0.338+0.542)+\exp(-0.247+0.430))} = 0.350$$

**Supplementary Results**

**Table S1.** Effect of being an initiator on the likelihood to signal during group departures. Estimated coefficients, standard error, and test statistics.

| | Estimate | Std. Error | z | P |
|---|---|---|---|---|
| Intercept | -1.433 | 0.082 | (1) | (1) |
| initiator | 1.774 | 0.189 | 9.389 | <0.001 |

(1) not meaningful in this context.

**Table S2.** Effect of signaling on the likelihood to initiate a group departure successfully. Estimated coefficients, standard error, and test statistics.

| | Estimate | Std. Error | z | P |
|---|---|---|---|---|
| Intercept | 1.715 | 0.362 | (1) | (1) |
| success | -0.031 | 0.472 | -0.066 | 0.947 |

(1) not meaningful in this context.

**Table S3.** Effect of belonging to the same unit on the interval times between dyads during group departures. Estimated coefficients, standard error, and test statistics.

| | Estimate | Std. Error | $\chi^2$ | Df | P |
|---|---|---|---|---|---|
| Intercept | 2.304 | 0.077 | (1) | (1) | (1) |
| belonging to same unit | -0.395 | 0.079 | -5.016 | 554.001 | <0.001 |

(1) not meaningful in this context.

**Table S4.** Estimated probabilities to progress in the front, middle, or rear positions of the file, in relation to age.

| | front | middle | rear |
|---|---|---|---|
| adult | 0.401 | 0.286 | 0.313 |
| young | 0.292 | 0.358 | 0.350 |

**Table S5.** Estimated probabilities to progress in the front, middle, or rear positions of the file, according to sex and male status.

| | front | middle | rear |
|---|---|---|---|
| female | 0.321 | 0.341 | 0.338 |
| primary male | 0.403 | 0.275 | 0.322 |
| non-primary male | 0.586 | 0.180 | 0.234 |

**Table S6.** Effect of being a female, a primary male, or a non-primary male on the likelihood of progressing in the first half of the front third of a group progression. Estimated coefficients, standard error, and test statistics.

| | Estimate | Std. Error | χ2 | Df | P |
|---|---|---|---|---|---|
| Intercept | -0.401 | 0.158 | (1) | (1) | (1) |
| primary male | 0.460 | 0.273 | 25.673 (2) | 2 | <0.001 (2) |
| non-primary male | 1.321 | 0.250 | (2) | (2) | (2) |

(1) not meaningful in this context;

(2) equal values because they refer to different terms of the same variable

**Table S7.** Effect of being a female, a primary male or a non-primary male on the likelihood of progressing in the first half of the rear third of a group progression. Estimated coefficients, standard error, and test statistics.

| | Estimate | Std. Error | $\chi^2$ | Df | P |
|---|---|---|---|---|---|
| Intercept | -0.196 | 0.112 | (1) | (1) | (1) |
| primary male | 0.227 | 0.211 | 17.388 (2) | 2 | <0.001 (2) |
| non-primary male | 0.976 | 0.230 | (2) | (2) | (2) |

(1) not meaningful in this context

(2) equal values because they refer to different terms of the same variable

**Table S8.** Effect of belonging to the same unit on the interval times between dyads during group progressions. Estimated coefficients, standard error, and test statistics.

| | Estimate | Std. Error | $\chi^2$ | Df | P |
|---|---|---|---|---|---|
| Intercept | 1.598 | 0.044 | (1) | (1) | (1) |
| belonging to same unit | -0.501 | 0.033 | -15.24 | 819.830 | <0.001 |

(1) not meaningful in this context.
